# Hedgehog signalling regulates patterning of the murine and human dentitions through Gas1 co-receptor function

**DOI:** 10.1101/2020.11.06.371476

**Authors:** Maisa Seppala, Beatrice Thivichon-Prince, Guilherme M. Xavier, Nina Shaffie, Indiya Sangani, Anahid A. Birjandi, Joshua Rooney, Jane N. S. Lau, Rajveer Dhalivar, Ornella Rossi, Adeel Riaz, Daniel Stonehouse-Smith, Yiran Wang, Laurent Viriot, Martyn T. Cobourne

## Abstract

The mammalian dentition exhibits wide numerical and morphological variation between different species. The regulation of dental pattern is achieved through complex reiterative molecular signalling interactions that occur through multiple stages of tooth development. We show that mice with loss-of-function in the Hedgehog co-receptor Gas1 have variation in size, morphology and number of teeth within the molar dentition. Specifically, premolar-like supernumerary teeth are present with high penetrance, arising through survival and continued development of vestigial tooth germs. We further demonstrate that Gas1 function in cranial neural crest cells is essential for the regulation of tooth number, acting to restrict Wnt signalling in vestigial tooth germs through facilitation of Shh signalling. Moreover, regulation of tooth number is independent of the additional Hedgehog co-receptors Cdon and Boc. Interestingly, further reduction of Shh pathway activity in a *Gas1* mutant background leads to fusion of the molar field and ultimately, developmental arrest of tooth development rather than exacerbating the supernumerary phenotype. Finally, we demonstrate defective coronal morphology in the molar dentition of human subjects carrying *GAS1* missense mutations, suggesting that regulation of Hedgehog signalling through GAS1 is also essential for normal patterning of the human dentition.

## INTRODUCTION

The mammalian dentition is a serially homologous structure defined by numerical and morphological variation in multiple species. The regulation of both tooth number and shape is achieved through complex molecular signalling interactions that occur reiteratively through multiple stages of tooth development to pattern the dentition (Cobourne and Sharpe, 2010; Jernvall and Thesleff, 2012; Tucker and Sharpe, 2004; Yu and Klein, 2020). Specifically, early signalling between the oral ectoderm and neural crest-derived (ecto)mesenchyme results in a thickening of ectoderm that develops into a rudimentary tooth bud. The bud stage tooth germ rapidly converts into cap and then bell stages to establish and refine coronal shape through further coordinated epithelial-mesenchymal signalling mediated by primary and secondary enamel knots, respectively (Jernvall and Thesleff, 2000; Jernvall and Thesleff, 2012) and representing a key component of morphogenesis. The molecular interactions that drive these stages of odontogenesis are dominated by activity of the Wnt, Bone Morphogenetic Protein (BMP), Fibroblast Growth Factor (FGF) and Hedgehog signalling families (Lan et al., 2014).

The ancestral mammalian dental formula consists of three incisor, one canine, four premolar and three molar teeth (O’Leary et al., 2013). The human dentition is therefore reduced, having lost an incisor and two premolar teeth during evolution; whilst the mouse dentition is highly reduced, consisting of only one incisor and three molars separated by an edentulous diastema region (Prochazka et al., 2010). Detailed analysis of the murine diastema has revealed the presence of paired vestigial tooth primordia that appear sequentially anterior to the first molar (M1) in both dental arches from embryonic day (E)13.5 (Peterková et al., 2002; Viriot et al., 2000). These primordia (termed R1, R2 and MS, R2 in the maxilla and mandible, respectively) disappear at the early bud stage through apoptosis (Peterkova et al., 2003); however, survival of R2 has been identified as the developmental basis of supernumerary premolar-like teeth that can occur in a number of mouse mutants (Ahn et al., 2010; Kassai et al., 2005; Klein et al., 2006; Mustonen et al., 2003; Ohazama et al., 2008; Peterkova et al., 2005; Zhang et al., 2003). Analysis of these mutants has demonstrated a complex network of molecular feedback inhibition that acts to restrict Wnt signalling and downstream FGF activity in R2, preventing developmental progression of this rudimentary tooth germ beyond the bud stage and therefore maintaining a reduced murine dental formula (Cobourne and Sharpe, 2010; Lan et al., 2014). Specifically, canonical Lrp5/6-dependent Wnt signalling induces the secreted Wnt-inhibitor Wise (also know as sostdc1, ectodin or USAG-1) in dental mesenchyme to establish a negative-feedback loop in R2 (Ahn et al., 2010); whilst FGF signalling is restricted in the epithelium and mesenchyme by activity of the Sprouty2 and Sprouty4 FGF-inhibitors, respectively (Klein et al., 2006). In addition, Sonic hedgehog (Shh) is a key target of Wise-mediated Wnt signalling, contributing to a Dkk1-dependent negative-feedback loop that acts to further inhibit Wnt activity in R2 (Ahn et al., 2010). The precise role of Shh within this regulatory model is not fully understood but a reaction-diffusion mechanism has been suggested, where a fine temporo-spatial balance between Wnt and Hedgehog signalling activity is responsible for dictating phenotypic outcome (Ahn et al., 2010; Cho et al., 2011).

Shh is a versatile developmental signalling molecule (Briscoe and Therond, 2013; Ingham and McMahon, 2001) that mediates pathway activity on the surface of receiving cells through ligand-binding to the principal Patched1 (Ptch1) receptor (Stone et al., 1996). Ptch1 ligand reception is facilitated by several proteins that act as co-receptors and includes the GPI-anchored membrane protein Gas1 (Growth arrest-specific1) (Allen et al., 2007; Martinelli and Fan, 2007) and the immunoglobulin superfamily member transmembrane proteins Cdon (cell adhesion associated, oncogene regulated) and Boc (Boc cell adhesion associated, oncogene regulated) (Kang et al., 1997; Kang et al., 2002). These co-receptors can bind Shh (Martinelli and Fan, 2007; McLellan et al., 2008; Okada et al., 2006; Tenzen et al., 2006), Ptch1 (Bae et al., 2011; Izzi et al., 2011) and in the case of Gas1, Ptch2 (Kim et al., 2020) directly on the surface of receiving cells and are collectively essential for vertebrate Hedgehog signaling (Allen et al., 2011). However, considerable redundency exists in co-receptor function between different tissues (Allen et al., 2007; Martinelli and Fan, 2007; Seppala et al., 2007; Tenzen et al., 2006), which may be reflective of differences in how these proteins influence binding of Shh-Ptch1 during signalling (Martinelli and Fan, 2007; McLellan et al., 2008). Moreover, *Gas1, Cdon* and *Boc* are all negatively regulated by Shh (Allen et al., 2007; Martinelli and Fan, 2007; Tenzen et al., 2006), placing them under complex transcriptional feedback control and further influencing the sensitivity of receiving cells to ligand (Petrov et al., 2017).

We have previously reported the presence of supernumerary premolar-like teeth in all jaw quadrants of mice lacking function of the Shh co-receptor Gas1 (Ohazama et al., 2008). Here we show that *Gas1*^*−/−*^ molars also have anomalies in cusp patterning and that the supernumerary teeth arise from survival and continued development of R2. We further demonstrate that *Gas1* function in cranial neural crest cells is essential for the regulation of tooth number, acting to restrict Wnt signalling in R2 through facilitation of Shh signalling. Interestingly, further reduction in Shh pathway activity in a *Gas1* mutant background does not exacerbate the supernumerary phenotype, rather leading to fusion of the molar field and ultimately, developmental arrest of tooth development. Finally, we demonstrate defective coronal morphology associated with the molar dentition of human subjects carrying missense mutations in *GAS1*, suggesting that regulation of SHH through GAS1 is also essential for normal patterning of the human dentition.

## RESULTS

### Multiple anomalies in the dentition of *Gas1*^*−/−*^ mice

We studied the arrangement and shape of maxillary and mandibular dental rows in *Gas1*^*+/−*^ and *Gas1*^*−/−*^ mice compared to wild type (WT) (Fig. 1A, B). In our sample, 80% of *Gas1*^*−/−*^ mice had at least one supernumerary tooth situated mesial to M1, more common in the maxilla compared to the mandible (46.3% versus 43.3%) and exhibiting significant morphological variation, ranging from mono to multi-cusped (particularly in the mandible). However, 55% of mutants had an absence of at least one M3 tooth, with 55.5% and 33.3% missing in the maxilla and mandible, respectively. In the maxilla, 50% of supernumerary teeth were associated with a missing M3 in the same row, whilst in the mandible this was 37.5%.

**Figure 1.**
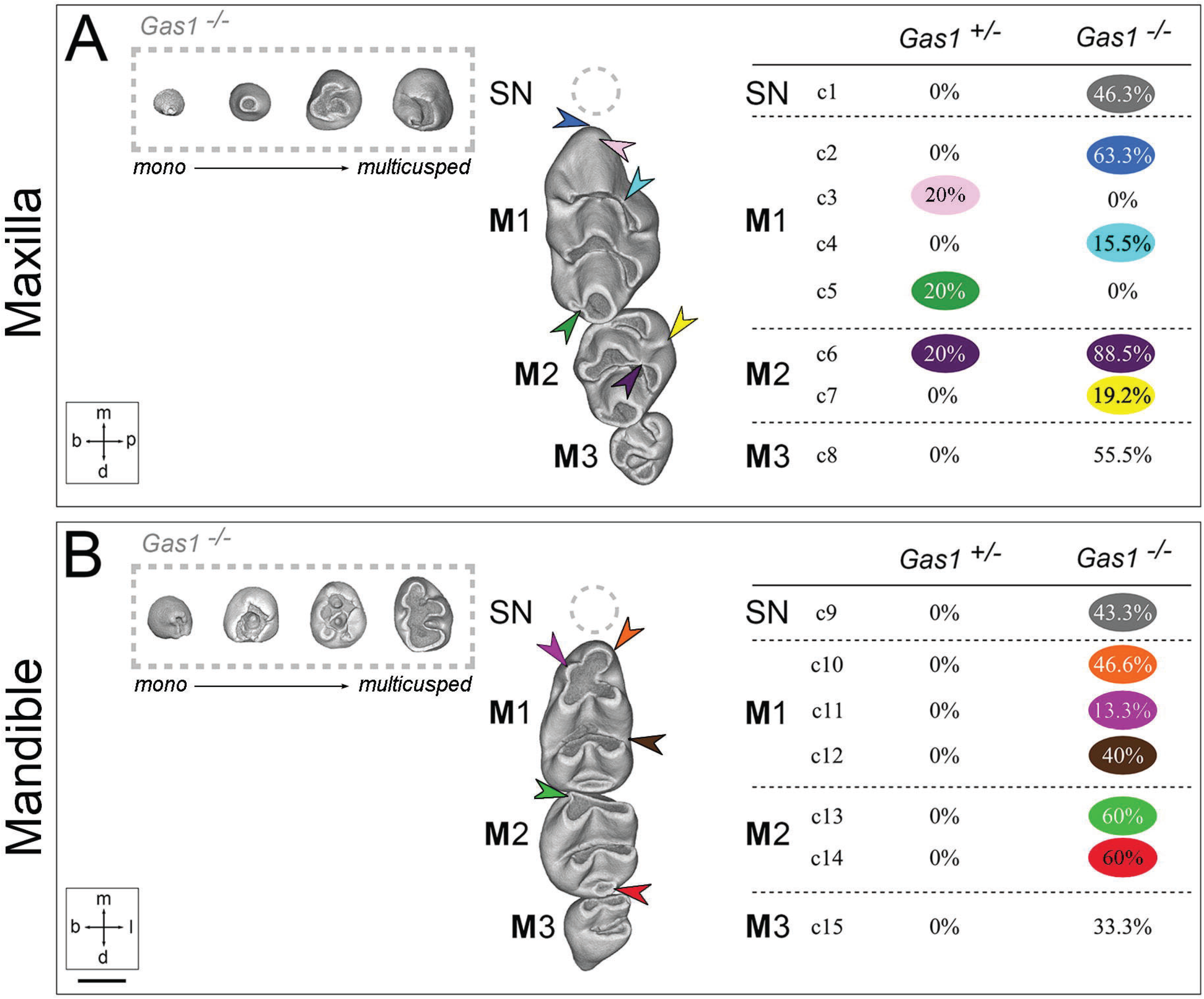
Dental character matrix. (**A**) Maxillary dental row (c1-8); (**B**) Mandibular dental row (c9-15). The first column displays WT maxilla (top) and mandible (bottom) molar tooth rows, with coloured arrowheads localising the most frequent defects identified in the mutant backgrounds. Each character is listed with the observed occurrence frequency in mutant mice. c1-8 are defects of the mutant maxillary molar dentition (c1, supernumerary tooth (grey hatched area); c2, straight M1 mesial cusp associated with a vertical tilt of the mesial component (dark blue arrow); c3, M1 extra mesial cusp (pink arrow); c4, absence of the M1 buccal cusp connection from the first chevron (light blue arrow); c5, connection pinching to disconnection of the M1 third chevron cusps (dark green arrow); c6, disconnection of the M2 mesio-lingual cusp from the first chevron (deep purple arrow); c7, abnormal connection between the two M2 lingual cusps (yellow arrow); c8, absence of M3). c9-15 are defects of the mutant mandibular molar dentition (c9, supernumerary tooth (grey hatched area); c10, absence of the M1 first lingual cusp (orange arrow); c11, absence of the M1 first buccal cusp (light purple arrow); c12, presence of an extra M1 first lingual cusp (brown arrow); c13, absence of the M3 first chevron cusp (light green arrow); c14, absence of the M2 most distal step (red arrow); c15, absence of M3). M1, M2, M3=first, second, third molar, respectively; SN=supernumerary tooth. m=mesial; d=distal; b=buccal; p=palatal (maxilla); l=lingual (mandible). Scale bar in B=0.45 mm for A, B.

All *Gas1*^*−/−*^ molar tooth rows also displayed anomalies in coronal morphology, with less severe variation also seen in heterozygous compared to WT teeth (Fig. 1A, B; Fig. 2A-C). In the maxilla of *Gas1*^*−/−*^ mice, the M1 mesial cusp was associated with a straight and vertically tilted additional component in 63.3% of teeth examined (a feature seen in all M1 associated with a supernumerary tooth) (Fig. 1A, Fig. 2C; dark blue arrows) and an absent palatal cusp connection from the first chevron in 15.5% (Fig. 1A, Fig. 2C; light blue arrows). In heterozygotes, an extra mesial cusp was present in 20% of M1 (Fig. 1A, Fig. 2B; pink arrows) with a connection anomaly present between both cusps of the upper M1 third chevron in 20% (Fig. 1A, Fig. 2B; dark green arrows). Both *Gas1*^*−/−*^ and heterozygous mice had a frequent disconnection of the M2 mesio-palatal cusp from the first chevron with a much higher frequency in the mutant (88.5% compared to 20%) (Fig. 1A, Fig. 2B, C; deep purple arrows). This anomaly was sometimes associated with an abnormal connection between the two palatal cusps of M2 in the mutant (Fig. 1A, Fig. 2C, yellow arrows). In the mandible, the heterozygous molar dentition was essentially normal (Fig. 1B, Fig. 2B) but a number of anomalies were identified in association with mutant teeth. Specifically, there was an absence of the mesio-lingual cusp in 46.6% of *Gas1*^*−/−*^ M1, with increased prevalence in the presence of a supernumerary (92.3%) (Fig. 1B, Fig. 2C; orange arrows); whilst the first buccal cusp was absent in 13.3% (observable in only 7.7% associated with a supernumerary) (Fig. 1B, Fig. 2B; light purple arrows) and an extra lingual cusp in 40% (Fig. 1B, Fig. 2B; brown arrows). In addition, 60% of *Gas1*^*−/−*^ M2 had an absence of the first chevron buccal cusp and the most distal cusp (Fig. 1B, Fig. 2B; light green and red arrows, respectively).

**Figure 2.**
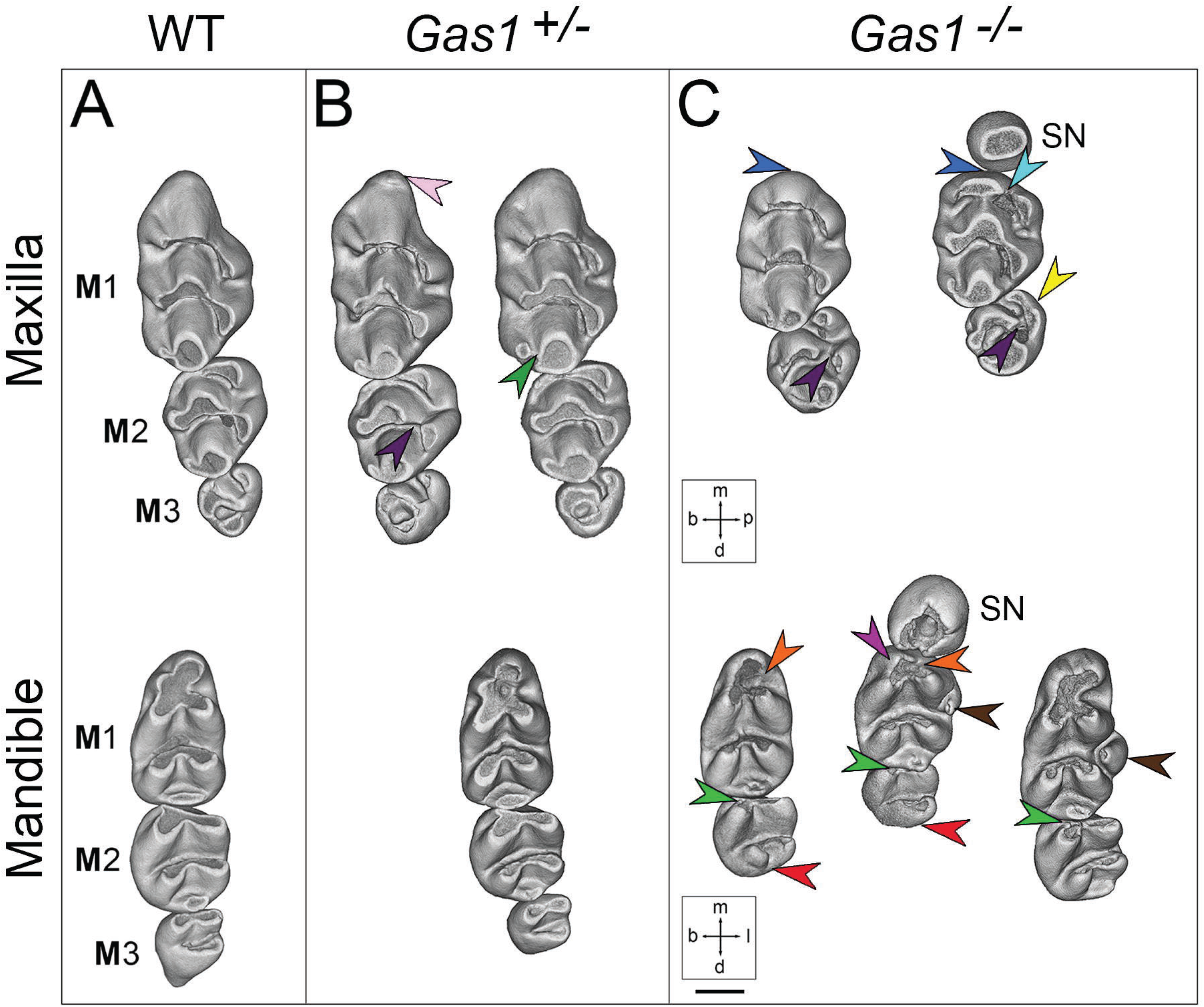
Abnormal phenotypes in the molar dentition of *Gas1*^*−/−*^ mice. (**A**) WT; (**B**) *Gas1*^*+/−*^; (**C**) *Gas1*^*−/−*^ dental row (maxilla above and mandible below). For the maxillary molar dentition: straight M1 mesial cusp associated with a vertical tilt of the mesial component (dark blue arrow); M1 extra mesial cusp (pink arrow); absence of the M1 palatal cusp connection from the first chevron (light blue arrow); connection pinching to disconnection of the M1 third chevron cusps (dark green arrow); disconnection of the M2 mesio-palatal cusp from the first chevron (deep purple arrow); abnormal connection between the two M2 palatal cusps (yellow arrow). For the mandibular molar dentition: absence of the M1 first lingual cusp (orange arrow); absence of the M1 first buccal cusp (light purple arrow); presence of an extra M1 first lingual cusp (brown arrow); absence of the M3 first chevron cusp (light green arrow); absence of the M2 most distal step (red arrow). M1, M2, M3=first, second, third molar, respectively; SN=supernumerary tooth. m=mesial; d=distal; b=buccal; p=palatal (maxilla); l=lingual (mandible). Scale bar in C=0.45 mm for A-C.

The identified morphological differences were also accompanied by size variation between the molar dentitions of WT and *Gas1*^*−/−*^ mice. In both the maxilla and mandible M1 length, width and tooth surface area was reduced in *Gas1*^*−/−*^ mice compared to heterozygote and WT (Fig. S1A-C). M1 dimensions in *Gas1*^*−/−*^ were 25% and 9% smaller than WT in the maxilla and mandible, respectively; whilst M2 was 34% and 37% smaller, respectively. The mean length of M1 was also decreased in both jaws and further reduced in the presence of a supernumerary. Whereas the *Gas1*^*−/−*^ M1 width was similar to heterozygote and WT in the maxilla; in the mandible, there was a significant increase in association with a supernumerary tooth or extra cusp. The M2/M1 area ratio in the mutant mandible was also significantly reduced compared to WT (Fig. S2). Interestingly, the number of cusps increased with tooth occlusal surface regardless of genotype. However, comparing WT and *Gas1*^*−/−*^ teeth demonstrated that very different surface values could contain the same number of cusps (Fig. S3A, B).

### Supernumerary teeth in *Gas1*^*−/−*^ mice are a product of the R2 vestigial tooth bud

We next investigated the developmental origins of supernumerary teeth in *Gas1*^*−/−*^ mice using three-dimensional (3D) reconstruction of serial histology (Fig. 3A-X). At E13.5, vestigial tooth primordia were visible in mesial regions of the maxillary and mandibular M1 bud stage tooth germs with no obvious gross morphological differences between WT and *Gas1*^*−/−*^ (Fig. 3A-D; E-H). At E14.5, the R2 vestigial primordia were still identifiable anterior to the cap stage M1 tooth germ in WT embryos, although some incorporation of R2 into the anterior region of the M1 cap was evident in the mandibular dentition (Prochazka et al., 2010; Viriot et al., 2000) (Fig. 3I, J; K, L). In *Gas1*^*−/−*^ mice, development of R2 continued toward an independent rudimentary cap stage in both arches, which was accompanied by some delay in formation of the M1 cap (Fig. 3M, N; O, P). At E15.5, the diastema buds were no longer distinct entities in the WT, particularly in the mandible where R2 was now a component of the anterior M1 cap (Fig. 3Q, R; S, T). However, clearly demarcated cap stage supernumerary tooth germs were present in *Gas1*^*−/−*^ embryos, anterior to a diminutive M1 cap stage tooth germ in both dental arches (Fig. 3U, V; W, X). These findings suggested that the developmental basis of supernumerary teeth observed in *Gas1*^*−/−*^ mice was prolonged survival of R2 vestigial tooth germs.

**Figure 3.**
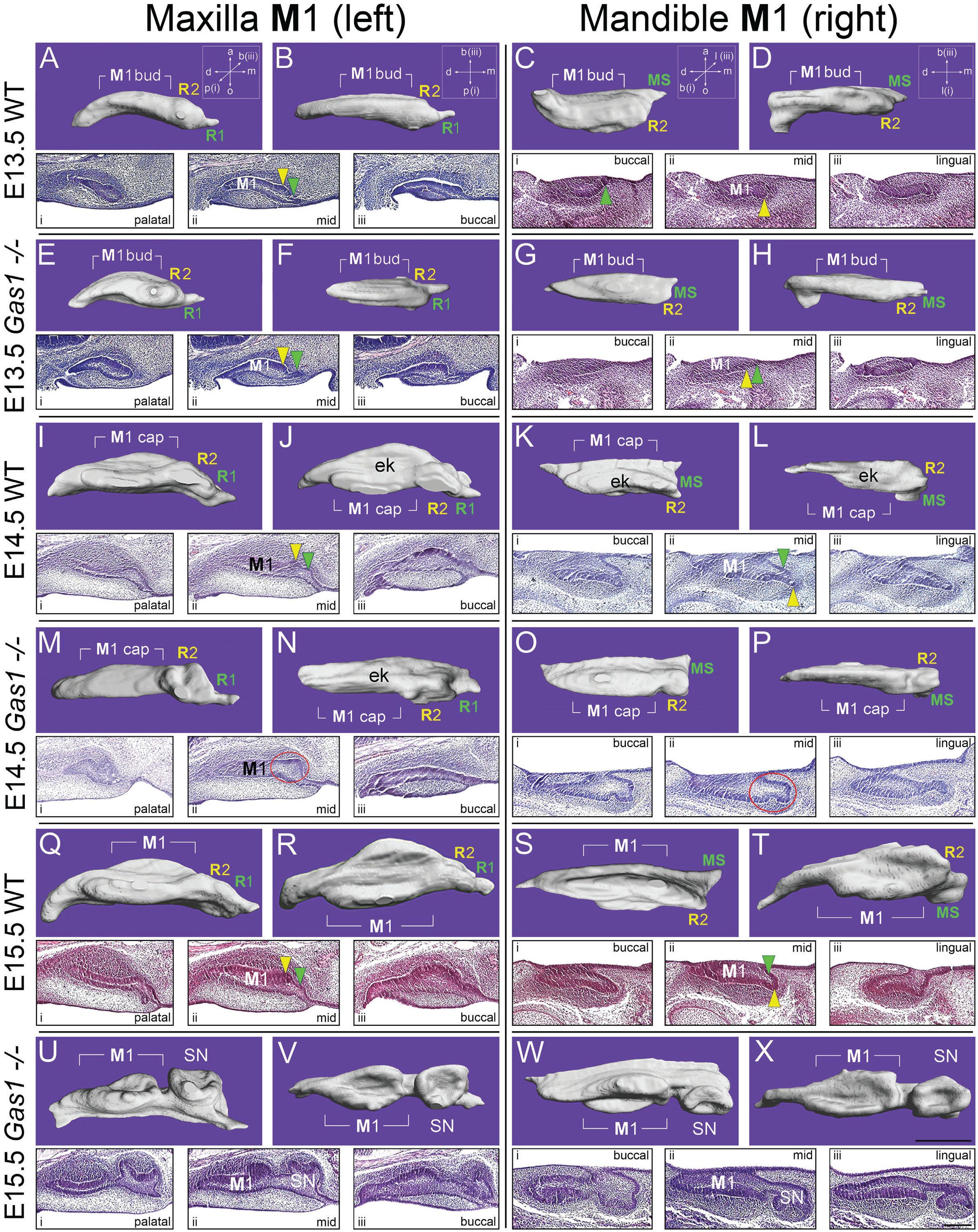
Supernumerary premolar teeth in *Gas1*^*−/−*^ mice are a product of the R2 vestigial tooth germ. 3D reconstructions of the epithelial component and serial para-sagittal histology of the left maxillary and right mandibular M1 in WT and *Gas1*^*−/−*^ mice. For the maxilla, (**A**, **E**, **I**, **M**, **Q**, **U**) shows 3D reconstruction of M1 from the palatal aspect; whilst (**B**, **F**, **J**, **N**, **R**, **V**) are orientated from below. For the mandible, (**C**, **G**, **K**, **O**, **S**, **W**) shows 3D reconstruction from the buccal aspect; whilst (**D**, **H**, **L**, **P**, **T**, **X**) are viewed from below. Serial histology is orientated from palatal through mid to buccal aspects (i, ii, iii; respectively) for the left maxillary M1 and from buccal through mid to lingual aspects (i, ii, iii; respectively) for the right mandibular M1. All images are orientated with mesial to the right. It can be seen that R1 and MS degenerate in the maxillary and mandibular M1 of WT and Gas1 mutant mice. However, whilst R2 degenerates in the WT maxillary and mandibular M1, in the Gas1 mutant this vestigial tooth bud survives and goes on to form a supernumerary premolar tooth in the maxilla and mandible. M1=first molar; SN=supernumerary tooth; ek=primary enamel knot. a=aboral; o=oral; m=mesial; d=distal; p=palatal; b=buccal; green arrowhead=MS (maxilla) and R1 (mandible); yellow arrowhead=R2 (maxilla and mandible); red circle highlights developing R2 in the mutant tooth germ. Scale bar in X=250 μm for A-X and in W-X iii (lingual)=250 μm for all histological sections.

It has previously been shown that molar patterning in the mouse dentition is a sequential process, which is initiated through the formation of vestigial tooth buds. This process has a characteristic molecular blueprint, which can be followed through the identification of *Shh* transcription in early signaling centres within the vestigial tooth buds and ultimately, the enamel knot of M1 at the cap stage (Prochazka et al., 2010). In order to confirm the contribution of vestigial tooth buds to supernumerary tooth formation in the absence of *Gas1* function at the molecular level, we undertook *in situ* hybridization for *Shh* on serial sections of the developing maxillary molar dentition and reconstructed these expression domains in 3D (Fig. 4A-F). In WT at E13.5, there was restricted *Shh* in R1 and a larger domain of expression in R2, whilst in *Gas1*^*−/−*^ the R1 *Shh* domain was more prominent and a clear R2 domain was also present. Significantly, the R2 region was identifiable at E13.5 in WT tooth germs as a localised region lacking proliferation and undergoing apoptosis; whilst in contrast, this region was actively proliferating and devoid of significant apoptosis in the mutant (Fig. S4A-D). At E14.5, the R2 domain had been lost in WT, whilst a large M1 domain was present in the enamel knot of the cap stage tooth germ. In the mutant at E14.5, there was evidence of prolonged R2 survival through continued *Shh* expression and delayed M1 development in more proximal regions, marked by an absence of *Shh* in the future enamel knot. In addition, there was a small region of ectopic *Shh* expression spanning the oral side of the epithelial cap (Fig. 4D, red arrows). At E15.5, in WT the M1 region of expression had begun to split into regions demarcating the secondary enamel knots, whilst in the mutant there was M1-associated *Shh* expression noted for the first time in the primary enamel knot and sustained expression in R2, which was now established as a cap stage supernumerary tooth germ. Collectively, these observations suggest that supernumerary premolar teeth observed in *Gas1*^*−/−*^ mice are a product of prolonged R2 survival and associated with delayed development of the mutant M1.

**Figure 4.**
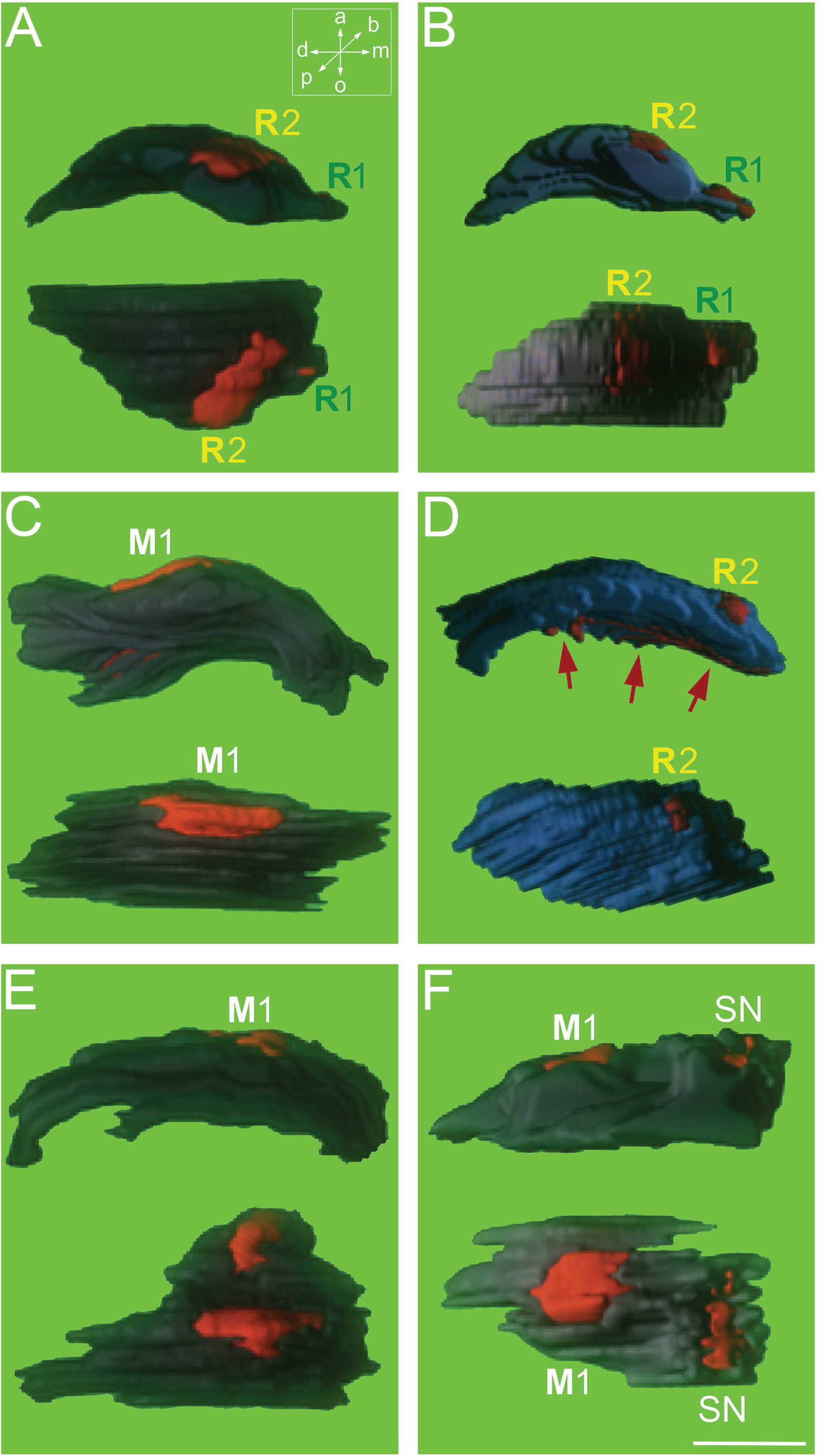
*Shh* expression during rudimentary tooth formation. 3D reconstructions of *Shh* expression (red) in the epithelial component (dark blue) of maxillary molar tooth germs derived from ^35^S radio-labelled *in situ* hybridization on para-sagittal sections. WT (left panels) and *Gas1*^*−/−*^ (right panels) at (**A**, **B**) E13.5, (**C**, **D**) E14.5 and (**E**, **F**) E15.5 showing palatal view above and aboral view below. *Shh* expression is seen to persist in R2 of the mutant tooth germ and the early supernumerary tooth with some ectopic activity also present in the aboral epithelium of the mutant tooth germ at E14.5 (red arrows in D). M1=first molar; SN=supernumerary tooth. a=aboral; o=oral; m=mesial; d=distal; p=palatal; b=buccal. Scale bar in F=250 μm for A-F.

### *Gas1* is expressed in odontogenic mesenchyme and epithelium during multiple stages of tooth development

In order to further investigate how loss of *Gas1* function influences tooth development, we undertook a detailed WT expression analysis in the developing molar dentition. Given that *Gas1* encodes a known co-receptor for Shh (Martinelli and Fan, 2007; Tenzen et al., 2006), we analysed expression within the context of Hedgehog pathway activity through comparison with *Shh* and *Ptch1* (Fig. 5A-O). At E12.5, *Shh* expression was visible in the maxillary and mandibular vestigial tooth buds R1 and MS, which was rapidly replaced by expression in R2 at E13.5 and ultimately, the primary enamel knot of M1 at E14.5 and early E15.5 (Fig. 5A-E). *Ptch1* expression demonstrated signalling activity localising progressively to the condensing mesenchymal cells contributing to the future dental papilla as well as the surrounding mesenchyme of the tooth germ from E12.5-15.5 (Fig. 5F-J). *Gas1* expression was initially predominantly reciprocal to *Ptch1* at E12.5 (Fig. 5K), reflecting the known negative regulation of *Gas1* by Shh in multiple regions of the embryo (Allen et al., 2007; Martinelli and Fan, 2007). At E13.5, *Gas1* had begun to localise within odontogenic mesenchyme condensing adjacent to the bud stage tooth germ (Fig. 5L) and by E14.5 there was strong expression in mesenchyme directly adjacent to the regressing R2 epithelial component of the tooth germ (Fig. 5M). However, by E15.5 the expression domain had changed dynamically, with transcripts now present in epithelium of the dental lamina and oral surface of the outer enamel epithelium (Fig. 5N arrowheads). In addition, the mesenchymal expression domain was now localised to a region situated between the oral epithelium and oral surface of the tooth germ, extending to posterior regions of the oral cavity on the buccal side (Fig. 5N, O arrowed). Collectively, these data were suggestive of a role for Gas1 in extending survival of the diastema tooth through the odontogenic mesenchyme but also potentially influencing epithelial function during morphogenesis of the enamel organ.

**Figure 5.**
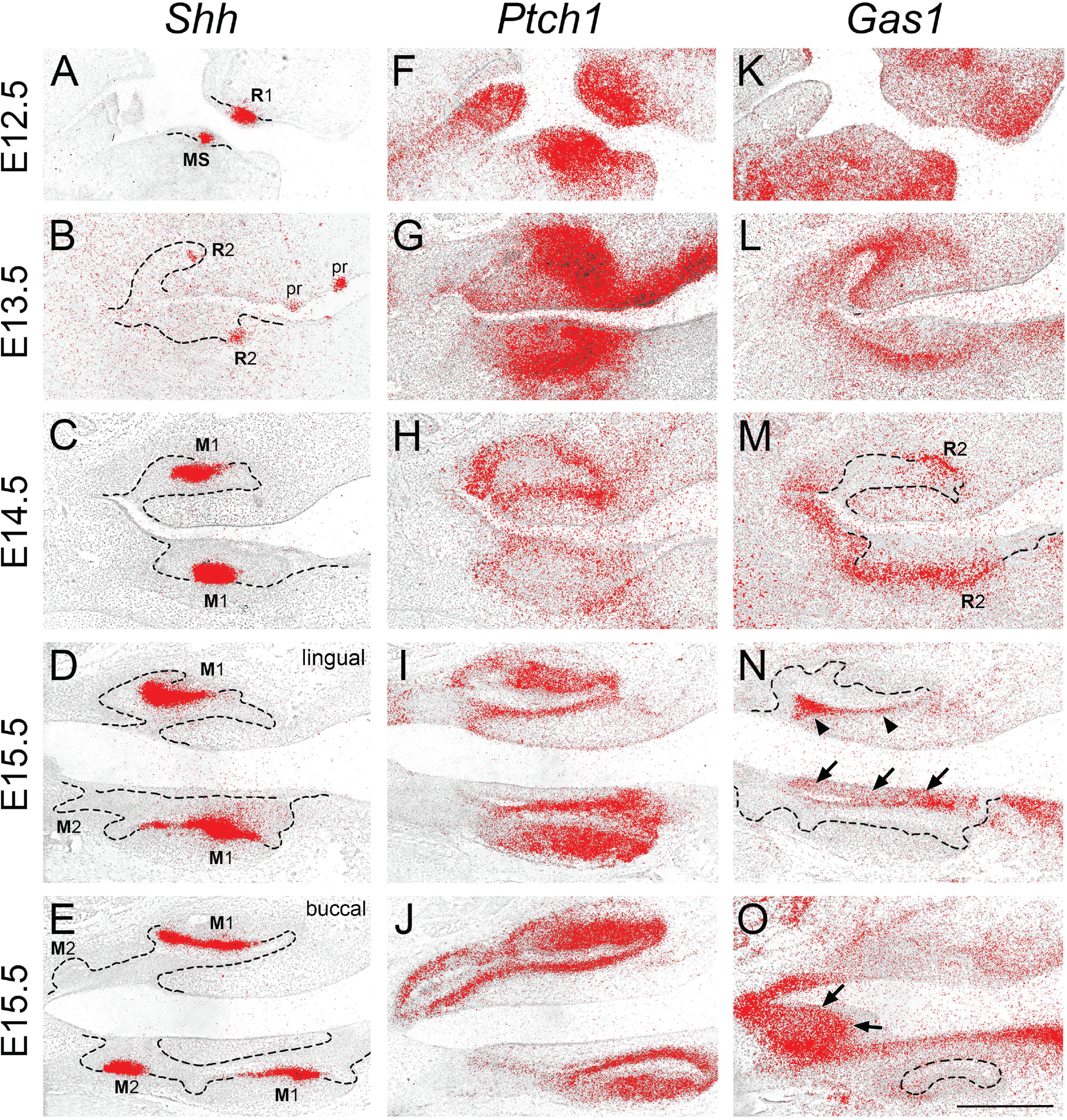
Shh pathway expression in the developing molar tooth germ. ^35^S radio-labelled *in situ* hybridization on para-sagittal sections through the developing maxillary and mandibular molar tooth germs from E12.5 to E15.5. (**A-E**) *Shh*; (**F-J**) *Ptch1*; (**K-O**) *Gas1*. Arrowheads in (**N**) indicate *Gas1* expression in the dental lamina and outer dental epithelium of the E15.5 tooth germ; arrows in (**O**) indicate *Gas1* expression in a mesenchymal domain extending to posterior regions of the oral cavity on the buccal side. Hatched black line represents epithelial boundary. Scale bar in O=500 μm for A-O.

### Gas1 regulates tooth number through odontogenic mesenchyme

Given the early expression of *Gas1* in odontogenic mesenchyme of the developing tooth germ we further investigated whether Gas1 function was essential within this tissue compartment for the regulation of tooth number. Specifically, we used *Wnt1-Cre* site-specific recombination to generate *Wnt1-Cre;Gas1*^fl/fl^ conditional mutant mice with ablation of *Gas1* function in cranial neural crest cells. These mice exhibited supernumerary premolar teeth situated mesial to M1 in all four jaw quadrants with a similar prevalence to that observed in *Gas1*^*−/−*^ mice. Thus, *Gas1* function is essential in murine odontogenic mesenchyme for the appropriate regulation of tooth number (Fig. 6A, B).

**Figure 6.**
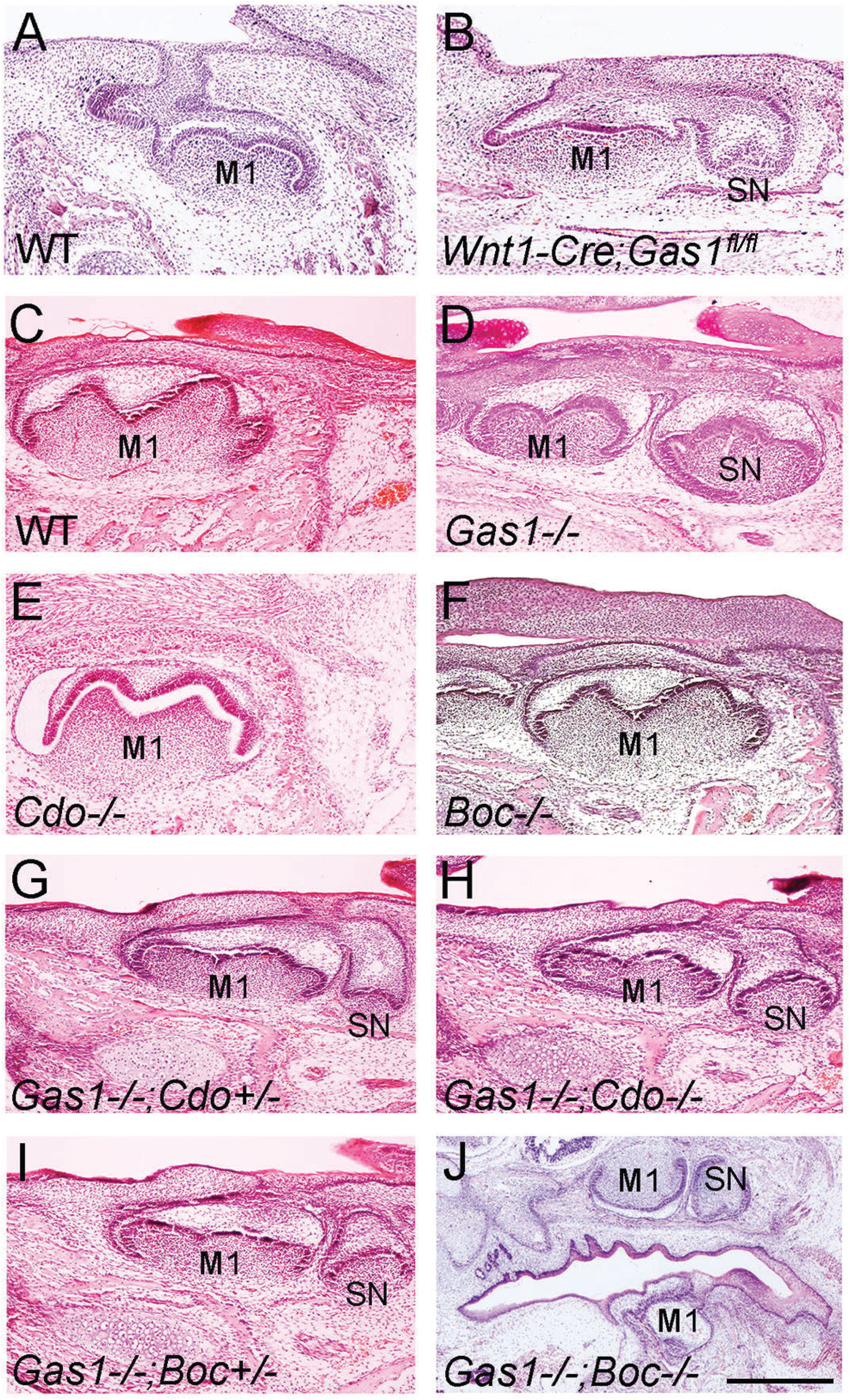
Molar phenotype of Hedgehog co-receptor mutant mice (**A**-**J**). Molar phenotype of E15.5 mice lacking Hedgehog co-receptor function. In comparison to WT (**A)**supernumerary premolar teeth a re present in *Wnt1-Cre;Gas1*^*fl/fl*^ conditional mice lacking Gas1 function in cranial neural crest cells.In comparison to (**C**) Wild type, (**D**) *Gas1*^*−/−*^ mice demonstrate supernumerary premolar teeth, whilst (**E**) *Cdo*^*−/−*^ and (**F**) *Boc*^*−/−*^ mice do not. (**G**) *Gas1*^*−/−*^;*Cdo*^*+/−*^, (**H**) *Gas1*^*−/−*^;*Cdo*^*−/−*^, (**I**) *Gas1*^*−/−*^;*Boc^+/−^* and (**J**) *Gas1*^*−/−*^;*Boc*^*−/−*^ all demonstrate the presence of premolar supernumerary teeth. Scale bar in H=500 μm for A-J.

### Hedgehog co-receptor function is partially redundant in the regulation of tooth number

Gas1 is known to interact with the Hedgehog co-receptors Cdon and Boc during signal reception in different developmental contexts and the combined loss of all three co-receptors results in an absence of pathway transduction (Allen et al., 2011). We investigated the expression domains of *Cdon* and *Boc* during murine tooth development and found transcripts in both the epithelial and mesenchymal components of the developing molar tooth germ between E12.5-15.5 (Fig. S5). Given that there are different tissue-specific requirements for these co-receptors throughout the developing embryo, including the craniofacial region (Allen et al., 2007; Cole and Krauss, 2003; Izzi et al., 2011; Martinelli and Fan, 2007; Okada et al., 2006; Sanchez-Arrones et al., 2013; Seppala et al., 2007; Seppala et al., 2014; Tenzen et al., 2006; Zhang et al., 2011; Zhang et al., 2006b), we investigated the effect of individual and combined loss of co-receptor function on tooth development. At E15.5, in contrast to *Gas1*^*−/−*^ mice, there was no evidence of supernumerary tooth formation in either *Cdon*^−/−^, *Boc*^−/−^ (Fig. 6C-F) or indeed, *Cdon*^−/−^;*Boc*^−/−^ double mutants (data not shown); whilst the loss of either single or both *Cdon* alleles in a *Gas1*^*−/−*^ background had no effect on prevalence of supernumerary tooth formation (Fig. 6G, H). *Gas1*^*−/−*^;*Boc*^*+/−*^ mice also displayed supernumerary premolars in all four jaw quadrants; however, *Gas1*^*−/−*^;*Boc*^*−/−*^ mice have lobar holoprosencephaly and rarely survive to E14.5 (Seppala et al., 2014), although we did find evidence of supernumerary premolar formation in the maxilla of one mutant (Fig. 7I-L).

**Figure 7.**
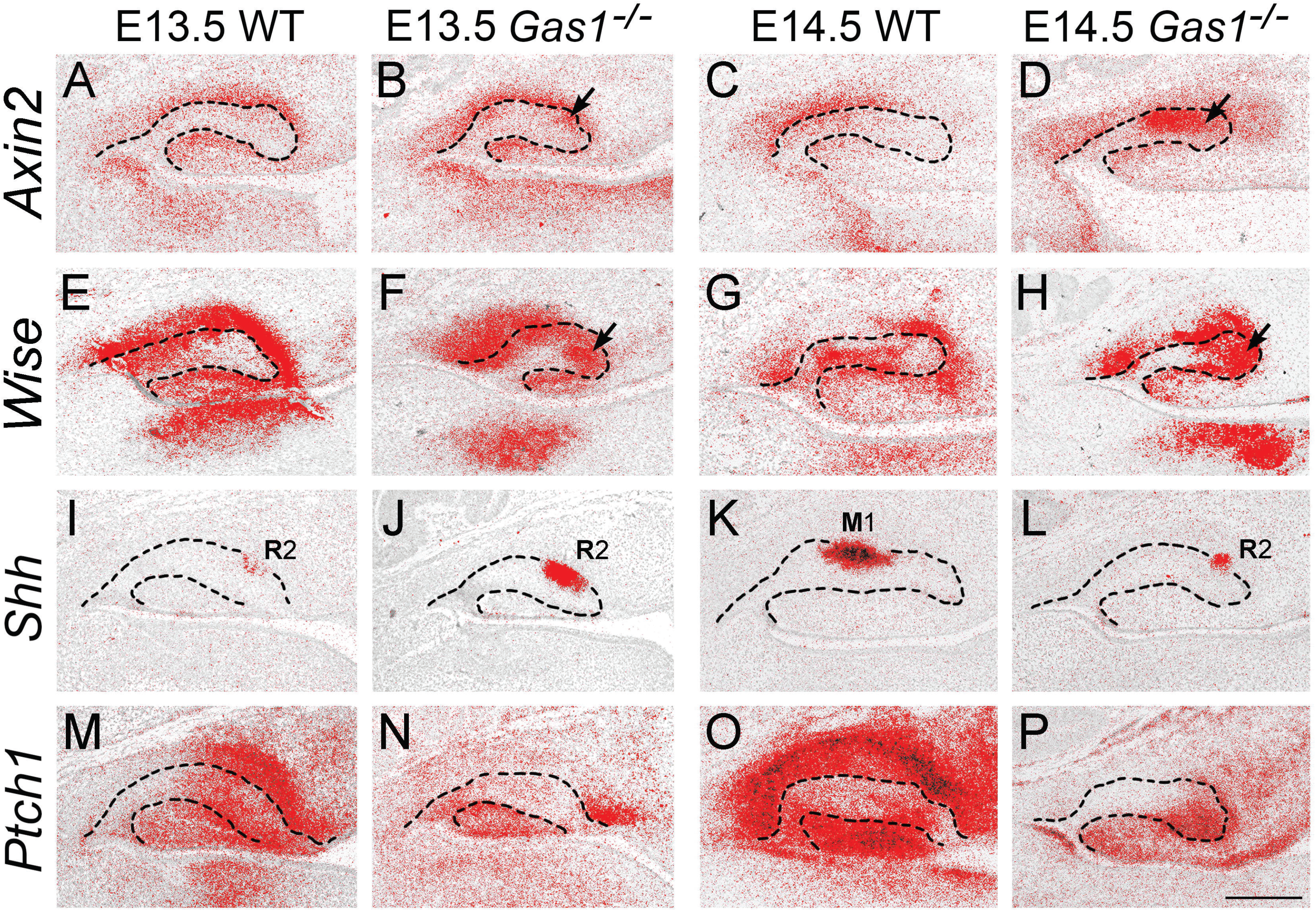
Increased WNT signaling and reduced Shh transduction in the developing R2 of *Gas1*^*−/−*^ mice. ^35^S radio-labelled *in situ* hybridization on para-sagittal sections through the developing maxillary molar tooth germs at E13.5 and E14.5. (**A**-**D**) *Axin2*; (**E**-**H**) *Wise*; (**I**-**L**) *Shh*; (**M-P**) *Ptch1*. At E13.5 and E14.5 there is increased expression of *Axin2* and *Wise* in epithelium and mesenchyme of the mutant R2 compared to WT (**A**, **B**; **C**, **D**and **E**, **F**; **G**, **H**) (black arrows indicate increased expression in R2). In contrast, despite sustained transcription of *Shh* in the mutant R2 (**I**, **J**; **K**, **L**) *Ptch1* expression was reduced (**M**, **N**; **O**, **P**) (black arrowheads). Hatched black line represents epithelial boundary. Black arrows indicate increased WNT signal transduction in R2. M1=first molar enamel knot. Scale bar in P=250 μm for A-P.

### Gas1 regulates tooth number through the Shh pathway

The level of Wnt signalling is carefully regulated in the WT R2 tooth bud with multiple negative feedback mechanisms in place to ensure that signal levels are maintained below the threshold required for survival and progression into a supernumerary tooth (Ahn et al., 2010; Klein et al., 2006). We investigated Wnt and Hedgehog signalling activity in WT and *Gas1*^*−/−*^ maxillary M1 tooth germs using section *in situ* hybridisation between E13.5 and E14.5 through analysis of the Wnt-target genes *Axin2* and *Wise* (Itasaki et al., 2003; Jho et al., 2002) and both *Shh* and *Ptch1*. At these stages in WT, *Axin2* was largely restricted to mesenchyme of the E13.5 and 14.5 tooth germs; whilst *Wise* expression was present in epithelium and mesenchyme of the WT (Fig. 7A, C; E, G). However, both these Wnt-targets were ectopically expressed in the persisting R2 epithelium of the mutant at both stages (Fig. 7B, D; F, H arrowed). Significantly, the increased Wnt activity observed in the mutant R2 was associated with reduced Shh signal transduction. Despite sustained and increased *Shh* transcription in the mutant R2 compared to WT at E13.5 and 14.5 (Fig. 7I-L), *Ptch1* transcription was reduced (Fig. M-P). An absence of Gas1 co-receptor function in mediating Shh signalling in odontogenic mesenchyme leads to increased Wnt signaling in R2.

A key component of these interactions is the influence of negative feedback between Wnt and Shh within a proposed reaction-diffusion system (Cho et al., 2011). Given the reduced levels of Shh transduction observed in R2 of *Gas1*^*−/−*^ mice we were interested in the consequences of further reducing signal levels in this mutant background. However, whilst *Gas1*^*+/−*^;*Shh*^*+/−*^ mice are phenotypically normal, loss of *Shh* alleles in a *Gas1*^*−/−*^ background leads to progressively more severe gross craniofacial defects. In *Gas1*^*−/−*^;*Shh*^*+/−*^ mice there are significant pattern defects affecting the posterior mandible and condylar region which cause gross disruption of molar tooth development (data not shown) (Seppala et al., 2007) whilst *Gas1*^*−/−*^;*Shh*^*−/−*^ mice only survive to E9.5. We therefore utilised the *Shh*^*tm6Amc*^ allele (*Shh*^*GFP*^), which encodes a bioactive Shh protein tagged with a green fluorescent protein (Shh::gfp) that is not processed as efficiently compared to WT (Chamberlain et al., 2008). *Shh*^*GFP/+*^ mice are normal, but *Shh*^*GFP/GFP*^ mice are hypomorphic, exhibiting embryonic lethality and significant phenotypic defects consistent with reduced Shh signal activity. These defects extended to the dentition where *Shh*^*GFP/GFP*^ mice have fusion between M1 and M2 but no supernumerary teeth (Fig. 7A, B) (Cho et al., 2011; Kim et al., 2019). The loss of a single *Gas1* allele did not affect the underlying molar phenotype in either a *Shh*^*GFP*^ heterozygous or homozygous background (Fig. 7C, D). In *Gas1*^*−/−*^;*Shh*^*GFP/+*^ mice, the gross craniofacial phenotype was more severe and we only identified supernumerary teeth in the maxilla (Fig. 7E). In contrast, *Gas1*^*−/−*^;*Shh*^*GFP/GFP*^ embryos failed to survive and were predominantly resorbed, with only one developing sufficiently to identify a lack of mandibular molars and evidence of only a poorly-formed molar tooth in the anterior maxilla (Fig. 7F). Abrogation of Shh function from E10.5 in *pCag-CreER™*;*Shh*^*fl/fl*^ mice resulted in molar development with severely disrupted morphology compared to WT (Fig. 7G, H).

Collectively, these findings demonstrate the importance of appropriately balanced Shh signal transduction during development of the molar dentition, with increased Wnt transduction in the absence of Shh inhibition leading to supernumerary tooth formation in *Gas1*^*−/−*^ mice. However, further reduction in Shh signal levels leads to more gross disruption of craniofacial and tooth development and an absence of individual supernumerary teeth.

### Loss of function in Hedgehog signalling affects human dental development

We further investigated the potential influence of Hedgehog signalling mediated through GAS1 function during development of the human dentition by examining the permanent dentition of three subjects previously identified with pathogenic mutations at the *GAS1* (n=2) or *GAS1* and *SHH* loci (n=1) and demonstrating features within the clinical spectrum of holoprosencephaly (HPE), comparing them to WT population-matched controls (n=3) (Ribeiro et al., 2010). The human M1 dental phenotype in these subjects was characterized by significant mesio-distal shortening and increased coronal width/length ratio (Fig. S6A-C). Moreover, there was some absence of specific cusps and modification of interconnections between cusps when compared to WT (Fig. 8A, B). Specifically, the shortened mandibular M1 had absence of the disto-buccal cusp (Hld, hypoconulid) in all subjects; whilst the maxillary M1 disto-palatal cusp (Hy, hypocone) was reduced or absent in one subject (*GAS1* c.775G>A). In the maxillary M1 of 2/3 subjects, a large and marked groove separated the disto-buccal cusp (Mc, metacone) from the mesio-palatal cusp (Pr, protocone), which removed the enamel bridge usually present between these two latter cusps. There was also evidence of shoveling affecting the maxillary incisor crowns in one subject, taurodontism in two subjects and generalised root-shortening. However, there was no evidence of any supernumerary teeth affecting the secondary (permanent) dentition (data not shown).

**Figure 8.**
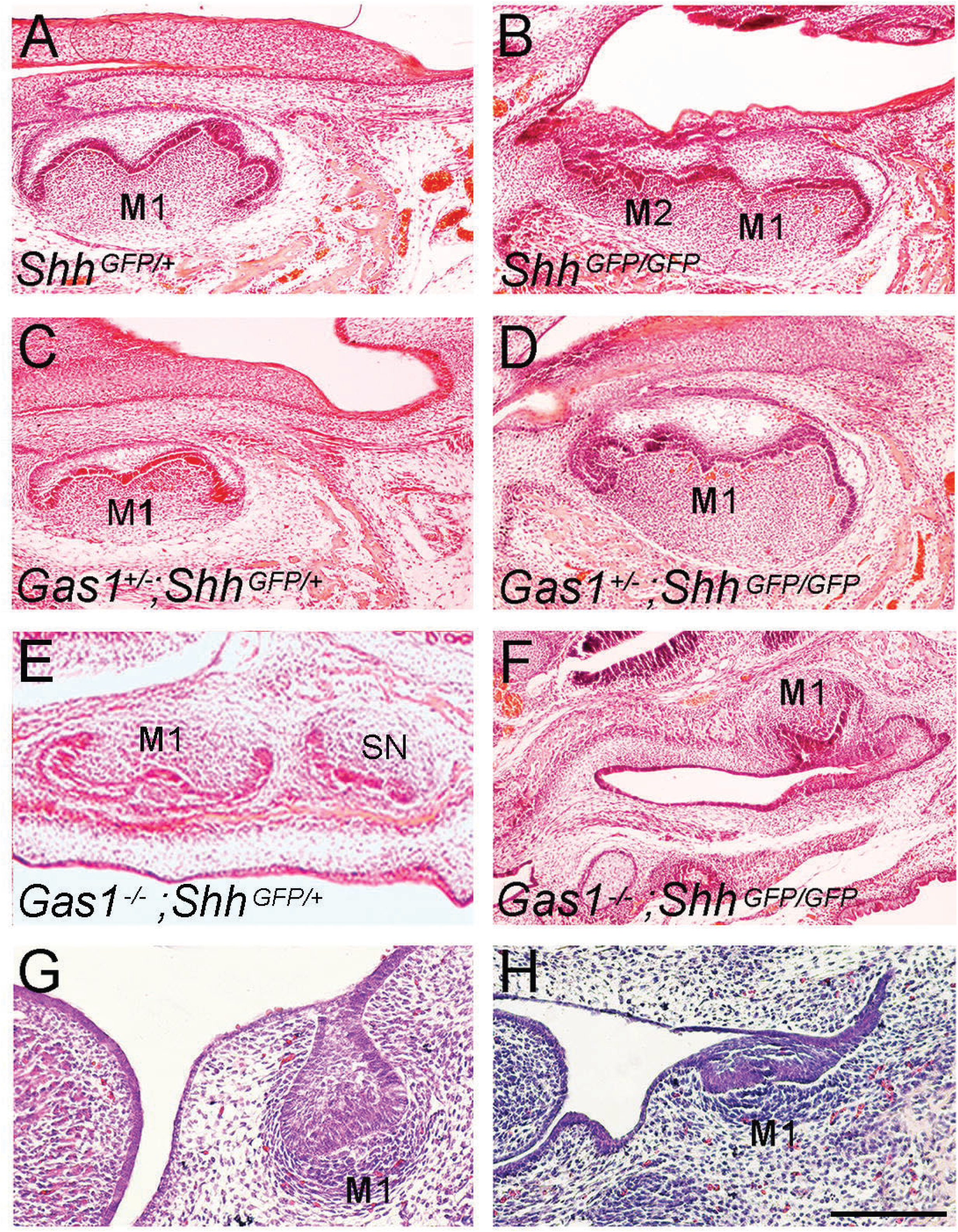
Spectrum of anomalies in the molar dentition of Shh pathway mutant mice. (**A**-**F**) Molar phenotype of E15.5 and (**G**, **H**) E14.5 mice lacking Shh pathway function. (**A**) *Shh*^*GFP/+*^, (**B**) *Shh*^*GFP/GFP*^, (**C**) *Gas1*^*+/−*^;*Shh*^*GFP/+*^, (**D**) *Gas1*^*+/−*^;*Shh*^*GFP/GFP*^, (**E**) *Gas1*^*−/−*^;*Shh*^*GFP/+*^, (**F**) *Gas1*^*−/−*^;*Shh*^*GFP/GFP*^ (**G**) WT, (**H**) *CreER™*;*Shh*^*fl/fl*^. Scale bar in H=500 μm for A-H.

**Figure 9.**
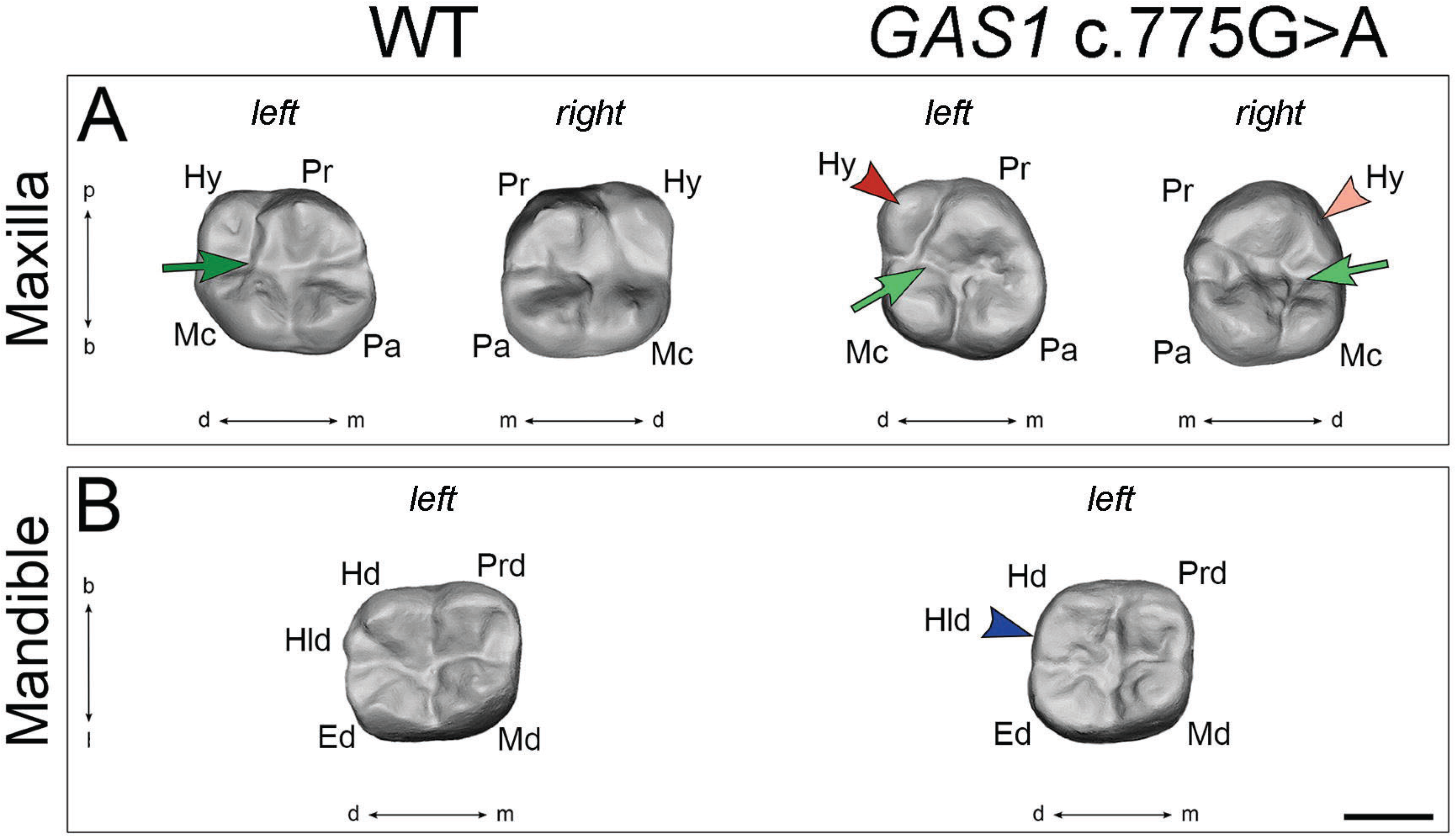
First molar phenotype of a human subject with *GAS1* c.775G>A missense mutation. (A) Maxillary (left and right) and (B) Mandibular (left) first permanent molars (M1) of representative population WT (left panel) and subject with *GAS1* c.775G>A missense mutation (right panel). In the WT maxillary M1, a bridge separated the metacone and protocone (dark green arrow) in the WT molar, which was replaced by a groove in the *GAS1* c.775G>A subject (light green arrows). In addition, the maxillary hypocone was reduced in size (red arrowhead) or absent (pink arrowhead). In the mandible, the *GAS1* c.775G>A M1 hypoconulid was absent (dark blue arrowhead). Ed, entoconid; Hd, hypoconid; Hld, hypoconulid; Hy, hypocone; Mc, metacone; Md, metaconid; Pa, paracone; Pr, protocone; Prd, protoconid; m=mesial; d=distal; b=buccal; p=palatal (maxilla); l=lingual (mandible). Scale bar in B=0.5 mm for A, B.

## DISCUSSION

In this investigation, we have identified morphological differences in the molar dentition of mice lacking function of the Hedgehog co-receptor Gas1, including variation in size and cusp pattern. These defects were seen in association with the formation of supernumerary premolar teeth and are consistent with a model of altered Shh signal transduction in the developing M1 tooth germ. Significantly, we also find evidence of molar cusp patterning defects in the first permanent molars of human subjects identified with loss-of-function mutations in *GAS1*. These findings suggest some conservation of developmental pathways in mice and humans during patterning of the dentition.

### Molecular interactions regulating cusp pattern

Shh has been shown to restrict cusp formation by regulating the spacing between cusps within individual teeth (Cho et al., 2011; Harjunmaa et al., 2012; Harjunmaa et al., 2014; Kim et al., 2019) and is also required for appropriate spatial patterning between teeth within the molar dentition (Cho et al., 2011). Specifically, a negative feedback reaction-diffusion model has been identified with Wnt signalling activating Shh in the enamel knot; which in turn, activates the Wnt-inhibitor Wise in surrounding mesenchyme (Cho et al., 2011; Kim et al., 2019). The effects of Shh inhibition on cusp patterning have been investigated in embryos following maternal administration of the Shh blocking antibody 5E1. The effects are variable, associated with both shape and locational diversity and influenced by multiple factors, including the timing of activity suppression (Cho et al., 2011; Kim et al., 2019). The maxillary M1 and mandibular M2 are the teeth predominantly affected with supernumerary cusps - located in the M1 mesial anterocone (prestyle), distal hypocone (posterocone) and mesial papracone, and M2 mesial metaconid (lingual anteroconid). *Gas1* heterozygous mice demonstrated an additional mesial cusp on the maxillary M1; whilst the mutant mandibular M1 was associated with an additional lingual cusp. However, the mutant also demonstrated absence of some cusps, including the mandibular M1 mesio-lingual cusp and M2 first chevron buccal cusp and most distal cusp. There may be a number of potential reasons for these variations; not least the fact that in order to study coronal anatomy in erupted teeth the *Gas1*^*−/−*^ mice analysed were those without cleft palate and surviving to adulthood, with a relatively milder craniofacial phenotype (Seppala et al., 2007). Moreover, the most severe molar pattern defects have been reported following pharmacological inhibition of Shh activity in the developing tooth from around E14.5 (Cho et al., 2011; Kim et al., 2019). In *Gas1*^*−/−*^ mice, whilst there is a reduction in Shh signalling within the tooth germ, transduction is not completely lost and therefore phenotypic changes might be expected to be more subtle, particularly within the context of a reaction-diffusion mechanism. In addition, *Gas1* is not expressed in the enamel knots but rather, in the external enamel epithelium and odontogenic mesenchyme at these stages. The presence of supernumerary teeth derived from R2 has also been shown to disrupt coronal morphology in a number of mouse mutants (Ahn et al., 2010; Kassai et al., 2005; Klein et al., 2006; Mustonen et al., 2003; Ohazama et al., 2008; Peterkova et al., 2005; Zhang et al., 2003), particularly in the mandibular M1 where R2 contributes to the mesial part of the crown during normal development (Peterkova et al., 2003). The variation in prevalence of supernumerary premolar teeth in these different developmental models is also likely to influence coronal anatomy of the resulting smaller M1. Interestingly, as the loss of Shh transduction increases or is instituted at progressively earlier stages of odontogenesis, the dental phenotype becomes more significantly associated with molar fusion and ultimately, a failure of appropriate invagination and growth of the tooth germ (Dassule et al., 2000; Gritli-Linde et al., 2002).

### Supernumerary tooth formation in *Gas1*^*−/−*^ mice

We have demonstrated that the developmental origin of supernumerary premolar teeth in *Gas1*^*−/−*^ mice is survival of the R2 vestigial tooth germ, consistent with a number of other mutant lines (Ahn et al., 2010; Kassai et al., 2005; Klein et al., 2006; Mustonen et al., 2003; Ohazama et al., 2008; Peterkova et al., 2005; Zhang et al., 2003). We also show that Gas1 function is required to facilitate Shh signal activity in odontogenic mesenchyme, further demonstrating the crucial role Shh plays in appropriate inhibition of Wnt signal transduction in R2, preventing progression of this vestige beyond a rudimentary bud stage. Wnt signalling acts upstream of FGF and Hedgehog pathways in the developing tooth germ (Ahn et al., 2010; Jarvinen et al., 2006; Kratochwil et al., 2002; Liu et al., 2008; Wang et al., 2009). In the absence of Gas1, Shh transduction is reduced in M1 mesenchyme and Wnt signalling in R2 epithelium elevates beyond a threshold required for supernumerary tooth formation. In this model, the relative levels of signal transduction within and around R2 are crucial, and therefore carefully regulated through multiple levels of negative feedback, involving Wnt, FGF and Hedgehog.

The importance of thresholds has been demonstrated through the analysis of *Wise* and *Shh*^*GFP/Cre*^ loss-of-function mutant mice, where individual heterozygotes exhibit normal tooth development; but double heterozygotes have supernumerary premolar teeth with high penetrance because the levels of Wnt signaling are increased sufficient to allow R2 to proceed to complete tooth formation (Ahn et al., 2010). In *Gas1*^*−/−*^ mice there is a similar molecular blueprint, Wnt signalling is increased beyond the required threshold because an absence of Gas1 reduces Shh signal transduction in odontogenic mesenchyme. However, the relationship with Shh signal transduction is complex because multiple levels of local feedback inhibition also exist within the pathway. The key targets of signal transduction *Ptch1*, *Ptch2* and *Hhip1* are all negative regulators of signalling (Chuang and McMahon, 1999; Marigo and Tabin, 1996; Motoyama et al., 1998); whilst the positive regulators *Gas1*, *Cdon* and *Boc* are all negatively regulated by pathway activation (Martinelli and Fan, 2007; Tenzen et al., 2006). These factors may further explain the subtle phenotypic variations seen in *Gas1*^*−/−*^ mice compared to those mutants for members of the Wnt and FGF pathways.

Interestingly, we found that *Shh*^*GFP/GFP*^ hypomorphic mice demonstrated fusions between M1 and M2, which is consistent with the phenotype associated with 5E1 inhibition from E14.5 (Cho et al., 2011; Kim et al., 2019). However, we did not observe any supernumerary premolar teeth in these mice, although their gross craniofacial phenotype was more severe. In *Gas1*^*−/−*^;*Shh*^*GFP/+*^ mice, the craniofacial defects are less severe but we only identified one supernumerary tooth in the maxilla with none in the mandible. In contrast, *Gas1*^*−/−*^;*Shh*^*GFP/GFP*^ mice had a severe form of HPE, lacked mandibular molars and only showed evidence of a poorly formed molar tooth in the anterior maxilla. A complete loss of signal from E10.5 in *pCag-CreER™*; *Shh*^*fl/fl*^ mice resulted in severely disrupted molar development consistent with *K14;Shh* and *Wnt1-Cre;Smo* loss of function mice (Dassule et al., 2000; Gritli-Linde et al., 2002).

It is likely that subtle differences in required levels of feedback inhibition contribute to variation in the penetrance of supernumerary tooth formation in different mutant lines. In *Gas1*^*−/−*^ mice with the most severe craniofacial defects including cleft palate, there is complete penetrance of supernumerary teeth in both jaws (Ohazama et al., 2009; Seppala et al., 2007), whilst in the mice analysed for coronal morphology reported here, who survive beyond birth the prevalence is around 50% in both arches. These figures compare with rates of around 60 per cent in both jaws for *Wise*^*−/−*^ mice and over 90 per cent and less than 10 per cent in the mandible and maxilla of *Sprouty2*^*−/−*^ mice, respectively (Klein et al., 2006).

### Hedgehog co-receptor function in tooth development

Shh signalling to odontogenic mesenchyme through the primary cilium plays a key role in the regulation of tooth number. In *Tg737*^*orpk*^ mice hypomorphic for the intraflagellar transport protein polaris, there is increased Shh transduction in odontogenic mesenchyme, which is associated with *Gas1* downregulation in this region and supernumerary premolar teeth with high penetrance (Ohazama et al., 2009). The precise details of how signal transduction is regulated through Hedgehog receptor function is not fully understood (Petrov et al., 2017; Xavier et al., 2016). However, we show here that despite being expressed in odontogenic mesenchyme *Cdon* and *Boc* are dispensable for the regulation of molar tooth number. Interestingly, neither Cdon or Boc have been identified at the primary cilium, which might explain their redundant role in the tooth. In addition, despite being collectively essential for Hedgehog pathway activity, only Gas1 seems to maintain a direct function in the promotion of signalling (Song et al., 2015). The individual and combined influence of *Gas1*, *Cdon* and *Boc* in craniofacial development has previously been shown to be variable and background-dependent in the mouse despite widespread expression (Allen et al., 2007; Cole and Krauss, 2003; Seppala et al., 2007; Seppala et al., 2014; Zhang et al., 2011; Zhang et al., 2006a). Our analysis suggests that in contrast to *Cdon* and *Boc*, *Gas1* plays the major role in regulation of murine tooth number.

### GAS1 function in human dental development

The human subjects identified with mutation in *GAS1* or *GAS1/SHH* had a craniofacial phenotype characterised by flat facial features, prominent eyes, maxillary and nasal hypoplasia, absent columella and bilateral cleft lip/palate. These features were seen in association with semilobar (*GAS1* c.599C>G) or mild HPE (*GAS1* c.775G>A); whilst the individual with a missense mutation in *SHH* (p.C363Y) and *GAS1* (c.863A>G) had a HPE-like phenotype (Ribeiro et al., 2010). There was a general reduction in length of M1 in these subjects and an absence of the hypoconulide. This is a relatively rare phenotype in modern human populations, representing around 0-10 per cent of individuals with a maximum frequency of 20 per cent in the populations of Western Eurasia. In addition, the hypoconulid is a cusp that appeared relatively late in the evolution of mammals, and is probably one of the last cusps to be established during development (Scott and Turner, 1997). The maxillary hypocone was absent in one subject with mild HPE. This is one of the four major cusps that compose the quadrangular upper molar and was the last major cusp addition to the M1 crown during the evolution of Primates. The absence of a hypocone associated with human M1 is therefore an extremely rare phenotype although this cusp may be reduced in size in some individuals (Scott and Turner, 1997).

## MATERIALS AND METHODS

### Mouse strains

All animal procedures were performed under approved licence (PPL7007441) in accordance with the Animal (Scientific Procedures) Act, Her Majesty’s Government, United Kingdom and local ethical approval at King’s College London and. *Gas1*^*+/−*^, *Cdon*^*+/−*^, *Boc*^*+/−*^, *Shh*^*tm6Amc/+*^, *Wnt1-Cre*^*+/−*^, Gas1^fl/+^, Shh^fl/+^ and pCag-CreER™ mice have been previously described (Chamberlain et al., 2008; Danielian et al., 1998; Dassule et al., 2000; Hayashi et al., 2003; Jin et al., 2015; Lee et al., 2001; Okada et al., 2006) and were used to generate single, compound and conditional mutants, respectively in a mixed 129sv/C57BL/6 background. For the analysis of erupted tooth morphology at 6 weeks old, *Gas1*^*+/−*^ mice in a 129sv/CD1 background were used to generate single mutants. Timed-matings were set up such that noon of the day on which vaginal plugs were detected was considered as embryonic day (E) 0.5. All tooth analyses involved a minimum of n=3 samples. In a mixed 129sv/C57BL/6 background *Gas1;Boc* compound heterozygous/mutant mice have compromised fertility and the yield of mutant embryos is significantly below that predicted by Mendelian ratios (Seppala et al., 2014).

### Imaging of dental tooth rows

The sample set for dental tooth row imaging consisted of 13 WT, 10 Gas1^+/−^ and 15 Gas1^−/−^ mice (a total of 152 dental rows or quadrants). The post-natal (P) specimen age ranged from 2 weeks to 1 year, which meant that M3 analysis was only possible for mice >3 weeks of age (M3 is only mineralised from P20) (Cohn, 1957). For each specimen, all tooth rows (left and right; upper and lower) were studied independently. All non-mineralized tissues were removed to allow observation of the dental rows. Tooth rows were imaged using conventional X-ray microtomography with a Pheonix Nanotom (General Electrics) using the following parameters: 70 kV tensions, 100mA intensity, 3000 images with time exposure of 500 ms and a voxel size of 3μm. 3D renderings were performed using VG Studiomax software (Volume Graphics, Germany). For each specimen, left and right, upper and lower rows were studied independently. Teeth were photographed using a Leica stereomicroscope with coronal length, width and occlusal area calculated using Leica Application Suite (LAS X) software (Leica Microsystems, Germany). Human maxillary and mandibular dental arches were imaged using 3Shape TRIOS scanners (3Shape, Denmark) from stone casts poured from maxillary and mandibular alignate impressions. Kruskal Wallis test was used to verify the significance of differences in tooth size. A threshold p value of 0.05 was used to assess the significance of the observed differences.

### Histological analysis and three-dimensional reconstruction of embryonic tooth germs

For histological analysis, embryos were fixed in 4% paraformaldehyde (PFA) at 4°C, dehydrated through a graded ethanol series, embedded in paraffin wax, sectioned at 7 μm and stained with haematoxylin and eosin. Continuous para-sagittal sections of the developing maxillary and mandibular molar regions were imaged and photographed using an Axioskop 2plus microscope, AxioCam HRC camera and Axiovision 3.0 software (Zeiss, Germany). Images were imported into Adobe Photoshop CS6 (Adobe, USA) software, edited to highlight the epithelial component of the developing tooth germs and saved. For *Shh* gene expression superimposition, continuous light-field images of ^35^S radioactive *in situ* hybridisation were taken and included with the relevant histology. Images were imported into DeltaViewer 2.1 3D Imaging software (freeware) and computationally stacked and aligned using the boundary of the oral epithelium and dental lamina as vertical and horizontal plane reference lines. In addition, continuous cross-referencing against the original histology was carried out throughout the process. The software reconstructed these images to produce a three-dimensional image of the developing molar tooth germs whilst excluding the background. The reconstructed surface was then smoothened and saved as a QuickTime 7.7.5 (Apple Corp, USA) movie file. Static images were also taken from different views for each developing tooth (top, oral, anterior, buccal and palatal views) and finally montaged in Adobe Photoshop CS6 to facilitate ease of comparison between control and mutant molars.

### *In situ* hybridisation

Section *in situ* hybridisation using ^35^S-UTP riboprobes was carried out as previously described (Gaete et al., 2015; Wilkinson, 1992). Light and dark-field images of sections were photographed using a Zeiss Axioscop microscope and merged in Adobe Photoshop CS6. We thank the following investigators for generously providing cDNA: Andrew McMahon, *Shh*; Matthew Scott, *Ptch1*; Chen-ming Fan, *Gas1*; Robert Krauss, *Cdon* and *Boc*; Atsushi Ohazama, *Wise* and *Axin2*.

### Proliferation and TUNEL assays

Bromodeoxyuridine (BrdU) labeling for cell proliferation was carried out on histological sections using a Zymed BrdU Labeling and Detection Kit (Invitrogen) according to the manufacturer’s instructions. Mouse embryos were labelled with BrdU via intra-peritoneal injection into pregnant females (5mg/100g body weight) 2 hours prior to sacrifice. Immunohistochemical detection of apoptotic cell death was carried out on histological sections using Terminal deoxynucleotidyl transferase-mediated deoxyUridine triphosphate Nick End Labeling (TUNEL). TUNEL was carried out using an APOPTag® Plus Fluorescein In Situ Apoptosis Detection Kit (Chemicon International) according to the manufacturer’s instructions.

### Analysis of human subjects

The analysis of human subjects was previously approved by the Institutional Review Board of the Hospital de Reabilitaeao de Anomalias Craniofaciais, Bauru, Brazil. Written informed consent was obtained from all parents and individuals included in the study. Human subjects with *GAS1* DNA sequence changes were identified from a sample of 54 individuals exhibiting features within the clinical spectrum of holoprosencephaly and a population-matched control sample with no history of familial structural anomalies of the central nervous system. Molecular analysis of genomic DNA identified the following variation in affected individuals: [Subject 1] was a female with variation c.775G>A in *GAS1;* [Subject 2] was a female with missense mutation c.808G>T in *GAS1* (and her father had a Leu218Pro mutation in *SHH*); [Subject 3]: a male with missense mutation c.863A>G in *GAS1* and p.C363Y missense mutation in *SHH*. The sequence analysis has been previously published with all identified *GAS1* variants considered to be damaging (Ribeiro et al., 2010). Dental panoramic radiographs and study casts were taken as part of routine dental care for these patients.

## ACKNOWLEDGEMENTS

We are indebted to the late Antonio Richieri-Costa, a geneticist at the Hospital for Rehabilitation of Craniofacial Anomalies, University of São Paulo who kindly provided the human dental records. He sadly passed away before the completion of this investigation. We thank Mathilde Bouchet for her help in the production of 3D reconstructions of dental rows using the Nanotom within the ANIRA-IMMOS platform of SFR Biosciences and Paul Tafforeau for his help in use of the Synchrotron X-ray microtomography at the ESRF in Grenoble. We are grateful to the following investigators for cDNA: Andrew McMahon, *Shh*; Matthew Scott, *Ptch1*; Chen-ming Fan, *Gas1*; Robert Krauss *Cdon, Boc*; Atsushi Ohazama *Wise, Axin2*. We thank Paul Sharpe and Cynthia Andoniadou for helpful comments relating to the manuscript.

## Supplementary Figures

**Supplementary Figure S1 Maxillary and mandibular M1 dental size in WT and *Gas1***^***−/−***^ **mice.** (**A**) M1 mesio-distal length (mm); (**B**) M1 buccal-lingual width (mm); (**C**) M1 coronal surface area (mm^2^). M1=first molar; blue solid diamond=*Gas1*^*−/−*^; blue outline diamond=*Gas1*^*−/−*^ with supernumerary premolar; orange diamond=*Gas1*^*−/−*^ with extra cusp; red square=*Gas1*^+/−^; green triangle=WT. * indicates significant difference (p<0.05).

**Supplementary Figure S2 Maxillary and mandibular M2 and M1 coronal surface ratios.** (A) M2/M1 coronal surface area ratios. M1=first molar; M2 second molar; dark blue diamond=*Gas1*^*−/−*^ maxilla; light blue diamond=*Gas1*^*−/−*^ mandible; dark green diamond=WT maxilla; light green triangle=WT mandible. * indicates significant difference (p<0.05).

**Supplementary Figure S3 Maxillary and mandibular M1 and M2 cusp number in WT and *Gas1***^***−/−***^ **mice.** (**A**) Maxilla; (**B**) Mandible. M1=first molar; blue solid diamond=*Gas1*^*−/−*^; red square=*Gas1*^+/−^; green triangle=WT.

**Supplementary Figure S4 Proliferation and apoptosis in the developing molar tooth germ of WT and *Gas1***^***−/−***^ **mice.** (**A, C**) BrdU and (**B, D**) TUNEL staining on sagittal sections through the developing maxillary M1 at E14.5. (**A, B**) WT; (**C, D**) *Gas1*^*−/−*^. Note the lack of BrdU-positive cells and presence of localised apoptosis in R2 of the WT (A, B; respectively) compared to the mutant (C, D; respectively). Scale bar in D=100 μm for A-D.

**Supplementary Figure S5 Shh receptor expression in the developing molar tooth germ.** Radioactive ^35^S *in situ* hybridization on frontal sections through the developing maxillary M1 from E12.5 to 15.5. (**A-D**) *Shh*; (**E-H**) *Ptch1*; (**I-L**) *Gas1*; (**M-p<**) *Cdon*; (**Q-T**) *Boc*. Scale bar in T=250 μm for A-T.

**Supplementary Figure S6 Maxillary and mandibular M1 dental size in human control subjects and those with *GAS1* mutation.** (**A**) M1 mesio-distal length (mm); (**B**) M1 buccal-lingual width (mm); (**C**) M1 width/length ratio. M1=first molar; green triangle=control subjects; blue solid diamond=subjects with *GAS1/SHH* missense mutation. *indicates significant difference (p<0.05).

